# Reassessing the contribution of the histone H3 tail to KRAB–DNMT3L-mediated epigenetic silencing

**DOI:** 10.64898/2026.09.24.753995

**Authors:** Minglei Song, Jiajun Lei, Jian-Kang Zhu

## Abstract

Neumann et al. introduced CHARM, a compact epigenetic silencer in which a histone H3 tail fused to DNMT3L was proposed to recruit and stimulate endogenous DNMT3A, enabling durable gene repression without a fused DNMT3A catalytic domain. Here, we evaluated the contribution of the H3 tail in independent reporter and endogenous-gene contexts. In an SNRPN reporter system, a KRAB–DNMT3L–dCas9 construct lacking the H3 tail displayed silencing kinetics comparable to CRISPRcharm Kv2, and mutating the critical H3K4 residue to alanine in CRISPRcharm Kv2 did not compromise this silencing. Similarly, after transient delivery of editor mRNAs to HEK293T cells, CRISPRcharm Kv2 did not consistently outperform the corresponding H3-tail-free construct at three endogenous loci, and mutating the critical H3K4 residue to alanine in CRISPRcharm Kv2 did not compromise this activity. These observations suggest that the engineered H3 tail does not confer a general functional advantage within the KRAB–DNMT3L–dCas9 architecture under the conditions tested.

## Main

Programmable epigenome editing enables targeted and potentially durable regulation of gene expression without altering the underlying DNA sequence. Combining transcriptional repressors with DNA methyltransferase-associated domains has enabled persistent gene silencing following transient editor expression (*1*). CRISPRoff subsequently established a programmable epigenetic memory platform by combining KRAB, the catalytic domain of DNMT3A (D3A), DNMT3L (D3L) and dCas9, achieving durable repression across a broad range of endogenous genes (*2*). However, the size of such multidomain editors presents challenges for delivery, motivating the development of compact architectures that exploit endogenous DNA methyltransferase activity.

CHARM (Coupled Histone tail for Autoinhibition Release of Methyltransferase) was developed to address this limitation by eliminating the fused DNMT3A catalytic domain (*3*). Its design builds on biochemical evidence that recognition of unmethylated histone H3 lysine 4 (H3K4me0) regulates de novo DNA methylation and that binding of the H3 tail to the DNMT3A ADD domain can relieve enzymatic autoinhibition (*4, 5*). In the original CHARM study, fusion of an H3K4me0-containing peptide markedly enhanced D3L–dCas9-mediated silencing in an endogenous CLTA reporter, whereas substitution of H3K4 with alanine abolished this enhancement. Subsequent incorporation of KRAB yielded optimized CHARM variants, including CRISPRcharm Kv2, which exhibited durable repression of endogenous cell-surface genes (*3*). However, KRAB–DNMT3L combinations have previously been shown to support persistent repression without a fused DNMT3A catalytic domain (*6*). This raises the question of whether the appended H3 tail provides an additional functional advantage once KRAB is incorporated into the DNMT3L-based editor.

To examine this question, we compared CRISPRoff, an H3-tail-free KRAB–D3L–dCas9 construct, CRISPRcharm Kv2, its H3K4A mutant and conventional KRAB–dCas9 CRISPRi **(Fig. 1A)**. We first evaluated their silencing kinetics using an independent SNRPN–EGFP reporter system. The four D3L-containing editors established sustained reporter repression over the 20-day observation period, whereas CRISPRi-mediated repression progressively declined after its initial peak **(Fig. 1B)**. Notably, CRISPRcharm Kv2 did not exhibit an evident improvement in silencing magnitude or persistence relative to the H3-tail-free KRAB–D3L–dCas9 construct. Furthermore, the H3K4A mutant retained a silencing trajectory closely resembling that of wild-type Kv2. Thus, the addition of the H3 tail did not confer a detectable silencing advantage, nor did mutation of its critical lysine 4 residue compromise editor efficacy in this reporter context.

**Fig. 1.**
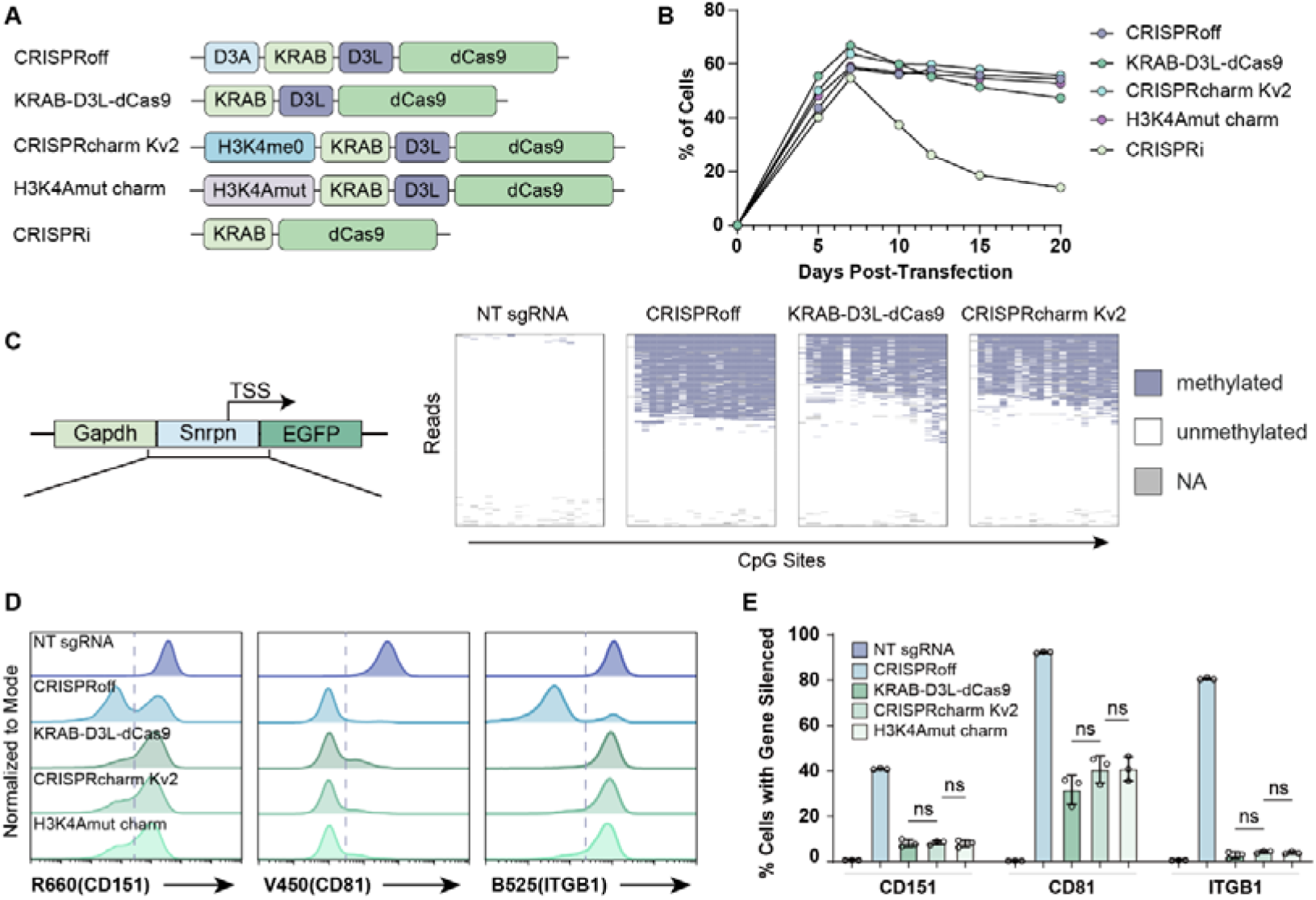
Comparison of CRISPRoff, KRAB–D3L–dCas9 and CRISPRcharm variants in reporter and endogenous-gene silencing assays. **(A)** Schematic representations of the five epigenetic editors: CRISPRoff, KRAB–D3L–dCas9, CRISPRcharm Kv2, CRISPRcharm Kv2-H3K4A and CRISPRi. D3A, DNMT3A catalytic domain; D3L, DNMT3L; H3K4me0, unmethylated histone H3 tail. **(B)** Time-course analysis of SNRPN–EGFP reporter silencing over 20 days following transfection with the indicated editors. The percentage of silenced cells was determined by flow cytometry. **(C)** Targeted bisulfite sequencing of the SNRPN reporter 20 days after transfection. The reporter architecture is illustrated on the left, and single-molecule CpG methylation patterns are shown for the indicated treatment groups. Rows represent individual sequencing reads, and columns represent CpG sites. Filled, white and gray positions indicate methylated, unmethylated and undetermined CpGs, respectively. **(D)** Representative flow-cytometry histograms of endogenous CD151, CD81 and ITGB1 (CD29) surface expression in HEK293T cells 10 days after electroporation with in vitro-transcribed (IVT) editor mRNAs and the corresponding sgRNAs. Dashed lines indicate the gating thresholds used to define silenced cells. **(E)** Quantification of CD151, CD81 and ITGB1 silencing by flow cytometry 20 days after IVT-mRNA electroporation. Bars represent mean ± SEM, with individual biological replicates shown as overlaid points. Statistical comparisons are indicated by brackets; ns, not significant.

To investigate whether sustained repression was accompanied by DNA methylation, we performed targeted bisulfite sequencing of the SNRPN reporter 20 days after transfection. Single-molecule analysis revealed extensive CpG methylation following CRISPRoff treatment and substantial methylation following treatment with either KRAB–D3L–dCas9 or CRISPRcharm Kv2, in contrast to the non-targeting control **(Fig. 1C)**. The two DNMT3A-catalytic-domain-free editors exhibited broadly similar methylation patterns, without an obvious additional methylation gain attributable to the H3 tail. These observations indicate that KRAB–D3L–dCas9 can establish targeted DNA methylation in the absence of the appended H3 peptide. As methylation of the H3K4A variant was not examined in this experiment, its retained silencing activity cannot be taken as direct evidence of unchanged methylation-writing activity.

We next extended the comparison to three endogenous genes, CD151, CD81 and ITGB1, in HEK293T cells. To examine editor performance under transient RNA delivery, we introduced in vitro-transcribed editor mRNAs together with the corresponding sgRNAs by electroporation. Flow-cytometric analysis at day 10 and quantitative assessment at day 20 revealed pronounced differences in the activity of the tested architectures **(Fig. 1D,E)**. CRISPRoff induced substantial repression of all three endogenous targets, whereas the DNMT3A-catalytic-domain-free editors exhibited marked locus dependence. At CD81, KRAB–D3L–dCas9, CRISPRcharm Kv2 and the H3K4A mutant each generated a substantial silenced population. By contrast, their activities at CD151 and ITGB1 were limited. Across the three loci, wild-type Kv2 conferred no consistent improvement over the H3-tail-free construct, nor did the H3K4A substitution noticeably diminish silencing efficacy, with none of the pairwise comparisons reaching statistical significance **(Fig. 1E)**.

Our observations are consistent with previous evidence that KRAB and DNMT3L can cooperate to establish persistent transcriptional repression without direct fusion of DNMT3A (*6*). They also highlight the context dependence of DNA methylation-based epigenetic silencing, as previously reported for other combinations of targeted epigenetic effectors (*7*). The substantially broader activity of CRISPRoff in our endogenous-gene experiments suggests that incorporation of the DNMT3A catalytic domain can overcome limitations encountered by the DNMT3L-based architectures under these conditions. However, the present data do not identify whether those limitations arise from endogenous methyltransferase recruitment, enzymatic activation or local chromatin properties.

Importantly, our experiments differ from those used to establish the original CHARM mechanism. We used an independent SNRPN reporter rather than the CLTA reporter and transient mRNA electroporation rather than the plasmid-based delivery employed in the original endogenous-gene experiments. Moreover, the original H3K4A loss-of-function observation was obtained in a KRAB-free architecture, whereas our comparison specifically addresses the optimized KRAB-containing Kv2 configuration. Our results therefore do not exclude a functional contribution of the H3 tail in other architectures or delivery contexts, nor do they challenge the established biochemical mechanism of H3K4me0-dependent DNMT3A activation.

In summary, our findings show that the engineered H3 tail is functionally dispensable within the optimized KRAB–DNMT3L–dCas9 architecture. They illustrate a critical gap between the biochemical capacity of an isolated peptide to stimulate an enzyme and its net phenotypic contribution when integrated into a complex epigenetic editor in living cells. More broadly, these results highlight that compact epigenome editor design requires rigorous benchmarking against true minimal baselines, ensuring that synthetic effector modules deliver genuine functional enhancements rather than architectural redundancy.

## Materials and Methods

### Plasmid construction

All epigenome editor plasmids (for direct transfection and for template IVT mRNA preparation) were generated using standard molecular cloning methods, following the architectural designs, domain sequences, and cloning strategies described by Neumann et al. (*3*). In brief, the CRISPRcharm Kv2 construct was assembled by fusing the unmethylated human histone H3 N-terminal tail to the KRAB–DNMT3L–dCas9 backbone, and its H3K4A variant was derived via site-directed mutagenesis. The minimal KRAB–DNMT3L–dCas9, CRISPRoff, and conventional KRAB–dCas9 (CRISPRi) constructs were modularly generated or sourced as previously reported (*2, 3*). Full annotated nucleotide and amino acid sequences for all expression and IVT template plasmids used in this study are provided in Supplementary Data 1.

### sgRNA design and preparation

All sgRNA spacer sequences targeting the *SNRPN* reporter and endogenous human loci (*CD151, CD81*, and *ITGB1*) were selected based on previously published studies (*2, 3*). Detailed spacer sequences and genomic target coordinates are listed in Supplementary Data 1. For transient plasmid transfections in reporter assays, sgRNAs were expressed under the control of the human U6 promoter. For electroporation-mediated endogenous gene editing, synthetic sgRNAs harboring standard chemical modifications were purchased from GenScript. A non-targeting (NT) sgRNA was used as a negative control across all experiments.

### In vitro transcription (IVT) of editor mRNAs

In vitro transcription of editor mRNAs was carried out as previously described (*8*). Briefly, linearized plasmid templates or PCR amplicons containing a T7 promoter were transcribed into mRNA incorporating modified nucleotides, capped, and purified according to established protocols. Synthesized mRNAs were quantified by spectrophotometry, and RNA integrity was confirmed by gel electrophoresis prior to electroporation.

### Cell culture and transfection

HEK293T cells and the clonal HEK293T SNRPN–EGFP reporter cell line were cultured in Dulbecco’s Modified Eagle Medium (DMEM, Gibco) supplemented with 10% (v/v) fetal bovine serum (FBS, Gibco) and 1% penicillin-streptomycin at 37°C in a humidified incubator with 5% CO□. Cells were routinely tested and confirmed negative for mycoplasma contamination.

#### Reporter assays (Plasmid transfection)

For the 20-day time-course experiment, SNRPN–EGFP reporter cells were seeded into 24-well plates at 3 × 10□ cells per well. After 24 hours, cells were co-transfected with 500 ng of editor plasmid and 500 ng of sgRNA plasmid using PEI according to the manufacturer’s instructions. Transfection medium was replaced after 24 hours.

#### Endogenous gene silencing (mRNA electroporation)

For endogenous locus targeting (*CD151, CD81, ITGB1*), parental HEK293T cells were harvested during logarithmic growth. A total of 1 × 10□ cells were resuspended in 100 μL of Neon Buffer R containing 5 μg of IVT editor mRNA and 5 μg of modified synthetic sgRNA. Electroporation was carried out using an Invitrogen Neon Transfection System (pulse voltage: 1250 V, pulse width: 20 ms, pulse number: 2). Immediately following electroporation, cells were rescued in pre-warmed complete growth medium and plated onto culture dishes. Cells were passaged every 2–3 days to maintain exponential growth throughout the 20-day monitoring period.

### Flow cytometry and cell surface immunostaining

Reporter EGFP fluorescence and endogenous cell-surface protein expression were evaluated at indicated time points (days 5, 7, 10, 12, 15, 20 for reporter kinetics; days 10 and 20 for endogenous genes) using a Beckman CytoFLEX flow cytometer:

#### EGFP reporter quantification

Transfected SNRPN–EGFP cells were dissociated with 0.05% Trypsin-EDTA, washed with PBS containing 2% FBS, and analyzed on the FITC/GFP channel (B525). Silenced cells were gated based on untransfected wild-type (GFP-negative) parental HEK293T controls.

#### Cell surface staining

For endogenous gene targets, cells were harvested at day 10 (representative histograms) and day 20 (quantitative silencing). Cells (1 × 10□) were washed and stained in the dark for 30 min at 4°C with fluorophore-conjugated primary antibodies: APC-conjugated anti-human CD151 (Clone 50-6, BioLegend, Cat# 350405, detected in R660 channel), Pacific Blue-conjugated anti-human CD81 (Clone 5A6, BioLegend, Cat# 349516, detected in V450 channel), and FITC-conjugated anti-human CD29 (Elabscience, Cat# E-AB-F0991C, detected in B525 channel). Gating thresholds for gene silencing were established using cells electroporated with non-targeting (NT) sgRNA. Flow cytometry data were analyzed using the floreada.io platform (https://floreada.io/).

### Targeted bisulfite sequencing and methylation analysis

Genomic DNA was isolated from SNRPN reporter cells 20 days post-transfection (NT sgRNA, CRISPRoff, KRAB–D3L–dCas9, and CRISPRcharm Kv2) using the FastPure Blood/Cell/Tissue/Bacteria DNA Isolation Mini Kit (Vazyme, Cat# DC112). Bisulfite conversion of genomic DNA was conducted using the EpiArt Ultrafast DNA Methylation Bisulfite Kit (Vazyme, Cat# EM112) according to the manufacturer’s protocol. The promoter-proximal region of the SNRPN reporter containing CpG sites was PCR-amplified using bisulfite-specific primers with 2× EpiArt HS Taq Master Mix (Vazyme, Cat# EM202). PCR amplicons were purified and submitted to Cwbiotech (Beijing, China) for targeted Oxford Nanopore sequencing at an average sequencing depth of ~1,000×. Single-molecule sequencing reads were aligned to the in silico bisulfite-converted reference sequence. Only full-length, high-quality reads covering all designated CpG positions were retained for single-molecule CpG methylation pattern visualization. Methylation status at each CpG site was categorized as methylated, unmethylated, or undetermined (NA).

### Statistical analysis

All data are presented as mean ± SEM of three independent biological replicates (*n* = 3), unless otherwise indicated. Statistical significance for multiple pairwise comparisons among the tested architectures at each endogenous locus was determined using one-way ANOVA followed by Bonferroni post hoc test calculated in GraphPad Prism (version 10.6). Differences were considered statistically significant at *P* < 0.05; ns denotes not significant (*P* ≥ 0.05).

## Competing Interests

The authors declare no competing interests.

## Author Contributions

M.S. and J.L. contributed equally to this work. M.S. raised the scientific question, constructed plasmids, and designed the study. J.L. conceived and designed the study, performed the experiments, and wrote the manuscript. J.-K.Z. acquired funding, supervised the project, and revised the manuscript. All authors discussed the results and approved the final version of the manuscript.

## Acknowledgements

This work was supported by the Guangdong S&T Program (2024B1111130001) and the Shenzhen Science and Technology Program (KJZD20240903102703005 and JCYJ20241202125311016).

## Notes

### Competing Interest Statement

The authors have declared no competing interest.

